# Transcriptome analysis of the human tibial nerve identifies sexually dimorphic expression of genes involved in pain, inflammation and neuro-immunity

**DOI:** 10.1101/450197

**Authors:** Pradipta Ray, Jawad Khan, Andi Wangzhou, Diana Tavares-Ferreira, Armen N. Akopian, Gregory Dussor, Theodore J. Price

## Abstract

Sex differences in gene expression are important contributors to normal physiology and mechanisms of disease. This is increasingly apparent in understanding and potentially treating chronic pain where molecular mechanisms driving sex differences in neuronal plasticity are giving new insight into why certain chronic pain disorders preferentially affect women versus men. Large transcriptomic resources are increasingly available and can be used to mine for sex differences and molecular insight using donor cohorts. We analyzed more than 250 human tibial nerve (hTN) transcriptomes from the GTex Consortium project to gain insight into sex-dependent gene expression in the peripheral nervous system (PNS). We discover 149 genes with sex differential expression. Many of the genes upregulated in men are associated with inflammation, and appear to be primarily expressed by glia or immune cells. In women, we find the differentially upregulated transcription factor SP4 that drives a regulatory program, and may impact sex differences in PNS physiology. Many of these 149 DE genes have some previous association with chronic pain but few of them have been explored thoroughly. Additionally, using clinical data in the GTex database, we identify a subset of differentially expressed (DE) genes in diseases associated with chronic pain, arthritis and type II diabetes. Our work identifies sexually dimorphic gene expression in the human PNS with implications for discovery of sex-specific pain mechanisms.

## Introduction

Sex-differential gene regulation and resultant changes in transcriptome, proteome and metabolome shape sexual dimorphic physiology and behavior in animals. Sex-differential molecular profiles in human tissues have a profound effect on health, resulting in disease susceptibility, prevalence and pathophysiology differences between sexes. Acute and chronic pain have a staggering global disease burden, and prevalence of many chronic pain conditions like fibromyalgia and neuralgia have been shown to be higher in women [1], and sex-differential molecular changes in peripheral (PNS) and central (CNS) nervous systems have been implicated in preclinical models [2]. Transcriptome profiles of human Dorsal Root Ganglia (hDRG) and Trigeminal Ganglia have been characterized previously [3, 4, 5] but these studies are underpowered for capturing subtle transcriptional changes between sexes. Human tibial nerve (hTN) transcriptomes, which contains axons from DRG neurons, along with a panel of other tissues harvested post-mortem, have been profiled using RNA-seq in hundreds of male and female donors as part of the GTEx project [6]. Some studies have characterized sex-differential gene expression changes [7, 8] and investigated the evolutionary and regulatory basis of such changes across the repertoire of tissues, but none have focused on the hTN. We cataloged sex differences in the hTN transcriptome focusing on potential functional impact in the context of pain, inflammation and neuro-immunity.

## Materials and Methods

### GTEx data requantification

PAXgene preserved hTN RNA-seq samples (dbGAP phs000424.v7.p2) with total RNA sequenced on the Illumina Truseq platform (and available donor consent information at the time of analysis) were identified. We mined the associated clinical information for samples to classify donors into 3 cohorts based on well understood phenotypic changes in peripheral nerves: those noted to have arthritis or rheumatoid arthritis (Chronic Joint Pain cohort, CJP), those noted to suffer from Type II diabetes (Type II Diabetes cohort, T2D, which causes diabetic neuropathic pain), and those without either of these diseases (baseline cohort, BSL). Samples noted to have sepsis, HIV infection, Type I diabetes or part of both CJP and T2D were not used. Statistical hypotheses testing to identify differentially expressed (DE) genes was performed on BSL (168:80 Male: Female Ratio / MFR) to characterize sex differences in healthy hTN (**Supp Table T1**), but hypotheses testing was not performed in CJP (11:10 MFR) or T2D (44:16 MFR) due to these cohorts being underpowered for DE gene identification (expression values, median fold change and Strictly Standardized Mean Differences for the two cohorts shown in **Supp Tables T2 and T3**). The GTEx uniform processing pipeline provided relative abundance of genes in the form of normalized read counts as Transcripts per Million (TPM). GTEx RNA-seq assays used rRNA-depleted total RNA libraries containing reads from non-polyA transcripts with the proportion of such reads potentially varying between samples [9]. We limited our analysis to validated coding genes by re-constraining the TPMs of coding genes (based on GENCODE annotation [10]) to sum to a million.

### DE gene identification

We performed nonparametric statistical hypotheses testing to identify DE genes (**Suppl Table T4**). To minimize effects of multiple testing, we filtered out lowly expressed or undetectable genes (based on the conservative filtering criterion of median gene TPM < 0.5 or maximum gene TPM < 1.0 in both males and females in a cohort). We also filtered out genes that were ubiquitously expressed in BSL by calculating the information theoretic score of normalized entropy (defined in Ray *et al* [4]) across all samples in a cohort. Since higher normalized entropy signifies more ubiquitous expression, we retained only those expressed genes in our analysis whose normalized entropy was less than the 75^th^ percentile of normalized entropies of expressed coding genes, thus performing gene filtering in a manner agnostic to the sex of the samples. Wilcoxon rank-sum test [11] was used to calculate p-values for differences in the male and female sub-cohorts in BSL for the median (50^th^ percentile) and the upper quartile (75^th^ percentile), which can be robustly estimated given the cohort size. To test for differences at the 50^th^ percentile (median), the entire male and female BSL sub-cohorts were used for comparison. In order to identify differences at the 75^th^ percentile (upper quartile), only values greater than or equal to the medians of the male and female BSL sub-cohorts were used. To account for multiple testing, the Benjamini – Hochberg Procedure (BHP) [12] was used for both the 50^th^ and 75^th^ percentile tests with a False Discovery Rate (FDR) threshold of 0.05 for both tests, (suggesting a combined FDR of 0.1). Since test statistics for the two tests are well correlated due to overlap of data, the empirical FDR is effectively in the range [0.05, 0.1].

## Results

We identified 29 genes in males, and 19 in females that were statistically significantly DE at 1.2-fold change or greater between the sexes in BSL at the 50^th^ percentile. Additionally, we identified an additional 74 genes in males (**Suppl figure S1**), and 27 genes in females that were DE at the 75^th^ percentile (**Suppl figure S2**). While several of these genes (including the 24 DE sex chromosomal genes) have been previously identified in the literature [7, 8], our analysis identified new DE genes including several that are potentially relevant to PNS function and pain mechanisms, including *ISLR2, SP4*, and *TPPP2*.

### Neural tissue enriched genes

We found several genes that are sex-differential in expression and are enriched in neural tissues (**Fig 1a**). To identify neural tissue enriched genes, we identified genes in our DE gene list with high (> 0.5) Neural Proportion Scores (quantified in Ray *et al* [4]) suggesting that they are primarily neuronally or glially expressed in CNS and PNS tissues. *NTRK1*, which is known to be expressed in mammalian sensory neurons, was upregulated in males at both the 50th and 75th percentiles, suggesting sex-differential axonal mRNA trafficking. *RNF165*, involved in axonal growth, was also upregulated in male samples at the 75^th^ percentile, and is another putative axonally transported mRNA. Well-known Schwann cell genes like Sonic Hedgehog (*SHH*) and Artemin (*ARTN*), that have been shown to be involved in pain, were also upregulated in males at the 75^th^ percentile. *ISLR2* and *MEGF11*, known to be involved in axonal pathfinding, were differentially upregulated in subsets of males and females respectively, and are potentially expressed in glia. Other neurally enriched genes that were differentially expressed in our datasets include *PGBD5* and RNA helicase *MOV10L1* in females (whose ortholog *MOV10* has been shown to be implicated in nerve injury response in rodents [13]), and *AGAP2, NXPH3* and *NRGN* in males. A small number of neural tissue enriched genes (*SHH, EFCAB5, FABP7*) showed stronger sex-dependent expression in the hTN compared to other tissues we examined (**Supp Table T5**).

**Figure 1.**
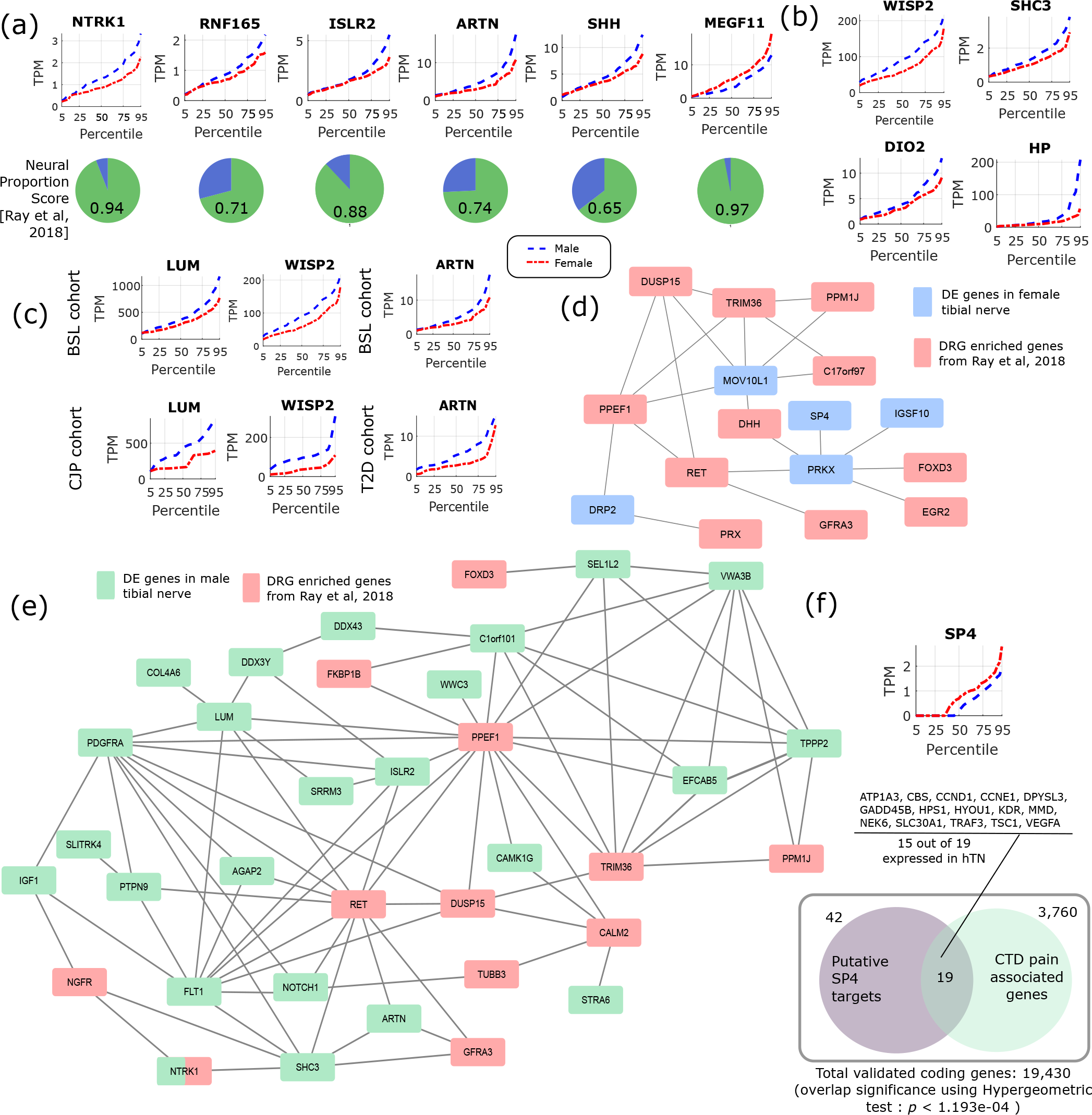
(a) Quantile plots for TPMs of sex-differentially expressed genes in BSL that are known to be preferentially expressed in neural tissue (having a Neural Proportion Score > 0.5 [4]). (b) Several examples of pro-inflammatory genes that are upregulated in male BSL sub-cohort. (c) *LUM* and *WISP2* show similar sex-differential expression trends in BSL and CJP, and *ARTN* shows similar trends in BSL and D2T. (d) Largest connected component of PIN based on StringDB interactions between mammalian DRG-enriched genes [4], and genes upregulated in female BSL sub-cohort. (e) Largest connected component of PIN based on known StringDB interactions between mammalian DRG-enriched genes [4], and genes upregulated in male BSL sub-cohort. (f) Quantile plot for *SP4* male and female sub-cohorts, and gene set enrichment analysis showing overlap of known *SP4* targets from TRANSFAC expressed in the hTN and Comparative Toxicology Database based pain associated genes.

### Pro-inflammatory gene signatures

The gene set upregulated in subsets of male BSL sub-cohort (**Figs 1a and b**) showed a different pro-inflammatory gene signature (*ARTN, HP, NOTCH1, CCL2, DIO2*, and others), with respect to the female BSL sub-cohort (*SULF1, GPR64, KRT4*), suggesting sex-differential expression patterns of inflammatory gene markers under normal conditions. This agrees with studies showing sex differences in clinical markers of inflammation [14]. Interestingly, the ratio of DE pro-inflammatory to anti-inflammatory genes in males were higher than in females, also in agreement with studies on human endothelial cells showing greater pro-inflammatory effects of androgens over estrogens [15]. We analyzed the BSL male sub-cohort pro-inflammatory genes in T2D and CJP, and found only 3 inflammatory genes (*LUM, WISP2, ARTN*) with sex-dimorphic expression at comparable effect sizes in either T2D or CJP with respect to BSL (**Fig 1c**) suggesting that the pro-inflammatory gene signature is unlikely to be caused solely by a subset of donors in BSL that are suffering from inflammatory conditions unreported in the clinical record. The larger number of genes upregulated in the male BSL sub-cohort can potentially be attributed to a combination of Y-chromosomal gene expression and downstream regulatory effects, and a larger set of upregulated pro-inflammatory genes.

### Potential Protein-Interaction Networks (PINs)

We investigated whether the gene products of the DE gene sets potentially interacted with genes whose expression is enriched in mammalian DRGs, which would help identify candidate sex-differential PINs in the PNS (**Figs 1d, 1e**). We used DRG-enriched genes from Ray *et al* [4] and our DE gene sets to identify putative PINs. The largest connected components from PINs generated using the StringDB database [16] show multiple DRG-enriched and sex-differentially expressed genes known to be expressed in the glia (*DUSP15, PRX, EGR2, DHH, FOXD3, ARTN*), and involved in pain and inflammation, which points to a potential role for glia in sex differential pain processing in human peripheral nerves (shown in preclinical models [17]). Additionally, the presence of several neuronally expressed genes in the interaction network among the set of DRG-enriched genes (*GFRA3, NTRK1, NGFR, RET* and *PPM1J*) also suggests sex-differential glia-neuron crosstalk, which in turn can affect neuronal plasticity and excitability differently between the sexes.

### DE regulatory genes

Underlying causes for such sex-differential gene expression include Y chromosome gene expression, incomplete X inactivation in females [7], differential androgen and estrogen receptor driven regulation between sexes [8] and transcription regulatory programs controlled directly or indirectly by sex chromosomal gene products. Additionally, we find differentially expressed transcriptional regulators on autosomes including *DMRTC1B* and *MED13* in males, and *SP4* in females. *SP4* is known to regulate pain-related genes, including several expressed in hTN (**Fig 1f**). While fold changes (between 1.2 and 1.5) in these autosomal genes under baseline conditions are unlikely to affect transcription in their regulatory targets, the presence of sex differential expression in a subset of samples suggests that under pathological conditions like inflammation and pain, more prominent sex-differential expression of these transcriptional regulators could potentially drive sex-specific regulatory programs.

### Roles in pain and neuro-immunity

Out of 149 DE genes we identified, 56 (>33%) genes have known roles in pain, inflammation and immune response (**Fig 2**). Sex differences in mammalian neuro-immune systems have been studied and linked to disease prevalence and susceptibility [18]. In hTN, we find that DE genes involved in the innate immune system (*C4A/B, CPAMD8* in males; *DDX3X, TF* in females) as well as genes in infiltrating or resident immune cells (*ADA, PDGFRA* in males; *CD2, CD8B* in females). We also found 34 genes that were associated with pain based on either the Comparative Toxicogenomics Database [19] or the Human Pain Genetics Database [20], including genes that are central to pain pathways like *NTRK1*, and *CCL2* in males and *TMEM97* in females [4, 21].

**Figure 2.**
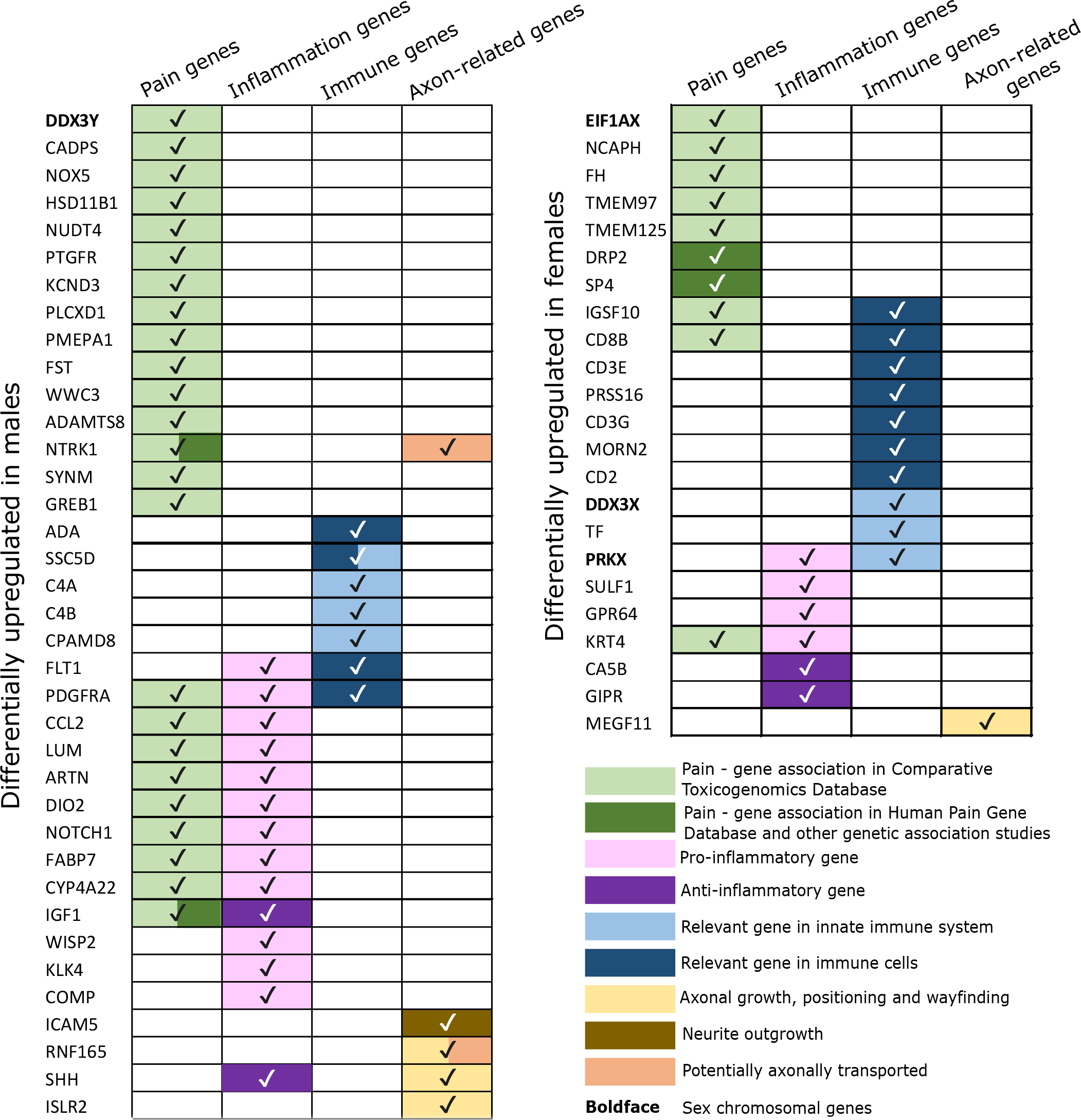
Annotations for sex differentially expressed genes in BSL showing whether they are involved in pain, inflammation, immune system functions, axonal growth or are potentially axonally transported, based on literature.

## Discussion

Our work provides a biostatistical framework, and thoroughly catalogs sex-differential gene expression in hTN. The public resource (https://www.utdallas.edu/bbs/painneurosciencelab/sensoryomics/sexdiffnerve/index.html) we generated provides a starting point for sex difference studies in human peripheral nerve drug target discovery, gene regulation and pathophysiology. We catalogued sex differences in gene expression in the PNS finding that many sexually dimorphic genes have known associations with pain, inflammation and neuro-immune interactions. This resource can be used for hypothesis-driven work to identify cell type and sub-compartment specificity of gene expression (e.g. *NTRK1, LUM*) by *in situ* hybridization or other techniques. It can also be used to back-translate into transgenic preclinical models to identify potential sex differences in mechanisms of PNS physiology that may be relevant to pain therapeutics (e.g. *SP4* as a key transcriptional regulator in the female PNS).

Our study can be expanded as GTEx cohort sizes continue to grow by identifying clinically relevant sex differences in CJP or T2D. Moreover, co-expressed (and putatively co-regulated) gene modules based on Whole Genome Correlation Network Analysis [22] can be identified by finding correlated expression changes across cohorts and sexes. Given that sex differences in gene expression likely contribute to sexual dimorphism in neurological disease, such as chronic pain, exploiting these transcriptomic resources will be increasingly important for mechanism and drug discovery.

## Author Contributions

PR, ANA, GD and TJP conceived the project. PR designed and performed experiments. JK, AW and DTF performed experiments. PR analyzed data. PR and TJP wrote the paper.

## Funding

NIH/NINDS grants R01NS065926 (to T.J.P.), R01NS102161 (to T.J.P. and A.N.A.) and R01NS098826 (to T.J.P. and G.D.)

## Conflict of Interest Statement

The authors declare there is no conflict of interest.

## Non-standard Acronyms

hTN: human Tibial Nerve
CJP: Chronic Joint Pain cohort
T2D: Type II Diabetes cohort
BSL: baseline cohort
DE: Differentially Expressed
BHP: Benjamini-Hochberg Procedure
MFR: Male:Female Ratio
PIN: Protein Interaction Network
DRG: dorsal root ganglion
PNS: peripheral nervous system

## Supplementary Material

**Supplementary Image S1** Differentially expressed male genes

**Supplementary Image S2** Differentially expressed female genes

**Supplementary Table T1** BSL male and female sub-cohort gene expression values

**Supplementary Table T2** CJP male and female sub-cohort gene expression values, fold changes and Strictly Standardized Mean Differences

**Supplementary Table T3** T2D male and female sub-cohort gene expression values, fold changes and Strictly Standardized Mean Differences

**Supplementary Table T4** BSL male versus female sub-cohort gene expression fold change and statistical hypothesis testing p-values, for statistically significantly changed genes

**Supplementary Table T5** Sex dimorphism scores across tissues

